# Enhanced targeted resequencing by optimizing the combination of enrichment technology and DNA fragment length

**DOI:** 10.1101/712125

**Authors:** Barbara Iadarola, Luciano Xumerle, Denise Lavezzari, Marta Paterno, Luca Marcolungo, Cristina Beltrami, Elisabetta Fortunati, Davide Mei, Annalisa Vetro, Renzo Guerrini, Elena Parrini, Marzia Rossato, Massimo Delledonne

**Affiliations:** Department of Biotechnology, University of Verona, Strada Le Grazie 15, 37134, Verona, Italy; Personal Genomics s.r.l, Via Roveggia 43B, 37136, Verona, Italy; Pediatric Neurology, Neurogenetics and Neurobiology Unit and Laboratories, Department of Neuroscience, A. Meyer Children's Hospital, University of Florence, viale Pieraccini 24, 50139 Florence, Italy

## Abstract

Whole-exome sequencing (WES) enrichment platforms are usually evaluated by measuring the depth of coverage at target regions. However, variants called in WES are reported in the variant call format (VCF) file, which is filtered by minimum site coverage and mapping quality. Therefore, genotypability (base calling calculated by combining depth of coverage with the confidence of read alignment) should be considered as a more informative parameter to assess the performance of WES. We found that the mapping quality of reads aligned to difficult target regions was improved by increasing the DNA fragment length well above the average exon size. We tested three different DNA fragment lengths using four major commercial WES platforms and found that longer DNA fragments achieved a higher percentage of callable bases in the target regions and thus improved the genotypability of many genes, including several associated with clinical phenotypes. DNA fragment size also affected the uniformity of coverage, which in turn influences genotypability, indicating that different platforms are optimized for different DNA fragment lengths. Finally, we found that although the depth of coverage continued to increase in line with the sequencing depth (overall number of reads), base calling reached saturation at a depth of coverage that depended on the enrichment platform and DNA fragment length. This confirmed that genotypability provides better estimates for the optimal sequencing depth of each fragment size/enrichment platform combination.

## Introduction

Next-generation sequencing (NGS) of targeted sets of regions of interest is one of the most widely used methods for genetic diagnostic testing [1–3]. Whole-exome sequencing (WES) platforms allow the enrichment of the entire set of human genes, offering diversity in terms of target region selection, bait length and density, the capture molecule, and the genomic fragmentation method [4, 5]. Despite some differences in the design of target regions, all current platforms perform well [7–11] thanks to improvements made over the past few years to enrich poorly-covered regions [5, 12]. The performance of WES is usually evaluated according to the depth and uniformity of coverage [6] because minimum site coverage of more than 10-fold [13–15] is generally required to identify germline variants [7]. However, current bioinformatics pipelines for the identification of variants generate a standard variant call format (VCF) file, which reports variant sites in genomes filtered by both site coverage and mapping quality [16]. In order to rescue variants in well-covered regions with a low mapping quality, base calling (genotypability) calculated by combining the confidence of read alignment with the depth of coverage should be considered as a more informative parameter for the assessment of WES performance [17].

There are many regions of low mapping quality in the human genome, often arising from repetitious sequences that prevent the unique alignment of short read pairs. Many genes have been duplicated over evolutionary timescales, and if the corresponding genomic regions are large enough to prevent a unique read alignment it becomes impossible to determine the source of each read [18]. Sequence aligners assign quality scores to read pairs according to the uniqueness of the alignment, so reads mapping to duplicated regions may gain a high quality score if one of the two read mates can be mapped unambiguously [18, 19].

WES library preparation protocols set the DNA fragment size to the average exon length, which is 170 bp in the human genome [20–22]. Short (< 100 bp) paired-end reads are generated to avoid the overlap of read pairs, but this fragment length is often shorter than duplicated regions. Furthermore, library preparation protocols often start from very low quantities of material (nanograms to picograms) [23], limiting the amount of DNA and consequently the number of unique fragments that can be produced. For this reason, 2 × 75 sequencing requires double the number of fragments to produce the expected depth of coverage that can be achieved by 2 × 150 sequencing. More amplification is therefore necessary, producing more PCR duplicates that must be removed during downstream data analysis, thus limiting the depth of coverage at target regions [24].

Here we describe a new approach that increases the standard DNA fragment size, allowing the longer fragments to extend beyond exonic regions to reach introns, which are under less selection pressure than protein coding sequences but still retain conserved polymorphisms [22]. We anticipated that such an approach would improve the mapping quality of DNA fragments in repetitious genomic regions.

## Materials and Methods

### Library preparation and exome capture

Samples NA12891 and NA12892 were obtained from the Coriell Institute for Medical Research, whereas sample VR00 was obtained from the whole blood of an unrelated third individual. Samples were processed using four different enrichment platforms: xGen Exome Research Panel V1 (IDT), SeqCap EZ MedExome (Roche), SureSelect Human All Exon V6 (Agilent), and the Human Core Exome Kit + RefSeq V1 (Twist). We produced three different DNA fragment lengths for each sample: short fragments based on the manufacturers’ recommendations (IDT = 150 bp, Roche, Agilent and Twist = 200 bp), medium fragments (expected length ∼350 bp), and long fragments (expected length ∼500 bp).

Libraries were prepared according to the manufacturers’ protocols. All DNA samples (IDT = 100 ng, Roche = 500 ng, Agilent = 1500 ng, and Twist = 50 ng) apart from Twist-200 bp, which were sheared enzymatically, were sheared using a Covaris M220 ultrasonicator, adjusting the treatment time to obtain the desired DNA fragment length (Supplementary Table 1). Given the low quantity of starting material for the Twist platform, the preparation of long DNA fragments was carried out twice for each replicate and the samples were combined before size selection to produce enough DNA to ensure sufficient library complexity. The size selection was performed with Agencourt AMPure XP (Beckman Coulter) before the pre-capture PCR. The DNA fragments and libraries were characterized using a Labchip GX Touch HS Kit (Perkin Elmer) or TapeStation (Agilent) to determine the size distribution and to check for adapter contamination.

Exome capture was performed independently for each combination of DNA fragment size and enrichment platform. Generally we followed the manufacturers’ protocols, but exceptions were made for the Agilent platform (single sample capture was performed) and for the preparation of long DNA fragments (the number of PCR cycles was increased by two for all platforms except Twist).

### Sequencing and bioinformatics

The samples were sequenced on a HiSeq3000 instrument (Illumina) in 75 bp paired-end mode for the short libraries and in 150 bp paired-end mode for the medium and long libraries. An in-house bioinformatics pipeline was developed for data analysis, integrating different software as described below.

Initial FASTQ files were quality controlled using FastQC (http://www.bioinformatics.babraham.ac.uk/projects/fastqc/). Low quality nucleotides have been trimmed using sickle v1.33 (https://github.com/najoshi/sickle) and adaptors were removed using scythe v0.991 (https://github.com/vsbuffalo/scythe). Reads were then aligned to the reference human genome sequence (GRCh38/hg38) using BWA-MEM v0.7.15 (https://arxiv.org/abs/1303.3997). The SAM output file was converted into a sorted BAM file using SAMtools, and the BAM files were processed by local realignment around insertion–deletion sites, duplicate marking and recalibration using Genome Analysis Toolkit v3.8 [16]. Overlapping regions of the BAM file were clipped using BamUtil v1.4.14 to avoid counting multiple reads representing the same fragment. Insert sizes were calculated after read alignment, measuring the distance of the two mates mapped on the genome using CollectInsertSize by Picard v2.17.10 (http://broadinstitute.github.io/picard/).

From the initial dataset representing each sample, we produced downsampled BAM files with a 140 theoretical X-fold coverage on the target design, subsampling the required number of fragments fragments (calculated as: (140 * design length) / (read length * 2)) using seqtk (https://github.com/lh3/seqtk). We then produced downsampled BAM files with a 10-80X-fold mapped coverage (the maximum mapped coverage value obtained by all the platforms, generated by sub-sampling the full dataset using sambamba v0.6.7 – https://github.com/biod/sambamba –).

We then used CallableLoci in GATK v3.8 to identify callable regions of the target (genotypability), with minimum read depths of 3 and 10. These values were integrated as additional WES performance parameters for the evaluation of variant detection. CollectHsMetrics by Picard v2.17.10 was used to calculate fold enrichment and FOLD 80 penalty values to determine enrichment quality. All WES performance parameters were calculated both on the design of each platform and on the standard dataset of RefSeq genes. For each sample, near target was defined as the distance from the region of interest corresponding to the average length of the DNA fragments. Variant calling was performed using GATK v4.1.2.

### Datasets

The RefSeq database (release 82) was downloaded from the UCSC Genome Table Browser (http://genome.ucsc.edu/). Online Mendelian Inheritance in Man (OMIM) genes associated with a clinical phenotype were downloaded from the OMIM website (https://www.omim.org/, release 15-05-2018).

## Results

### Influence of DNA fragment size on duplicate and off-target rates

We assessed the performance of short (∼200 bp), medium (∼350 bp) and long (∼500 bp) DNA fragments on four major commercial exome enrichment platforms produced by IDT, Roche, Agilent and Twist (Table 1). For each platform, libraries were generated from the genomic DNA of three unrelated individuals and were enriched according to the manufacturers’ instructions, and then sequenced on an Illumina HiSeq3000 instrument. Short libraries were sequenced in the 2 × 75 bp format, whereas medium and long libraries were sequenced in the 2 × 150 bp format.

**Table 1.** The 140X dataset. For each platform and DNA fragment length combination, the 140 theoretical X-fold coverage is shown for the target design dataset (mean of the three independent experiments). The columns show the average insert size, percentage of reads marked as duplicates, mapped coverage on the target, percentage of the target covered by at least 1, 5, 10, 20 and 30 reads, percentage of callable bases on the target for standard read depth (>3) and read depth >10, percentage of bases on/near/off target, fold enrichment and FOLD 80 base penalty. DNA fragment lengths: S = short, M = medium, L = long.

| ID | Average insert size | % Duplicates | Mapped coverage (X) | %1X | %5X | %10X | %20X | %30X | % PASS | % PASS RD>10 | % ON TARGET | % NEAR TARGET | % OFF TARGET | Fold enrichment | FOLD 80 penalty |
| --- | --- | --- | --- | --- | --- | --- | --- | --- | --- | --- | --- | --- | --- | --- | --- |
| IDT-S | 171.61 | 12.61 | 69.38 | 99.82 | 99.77 | 99.61 | 98.14 | 92.96 | 96.81 | 96.66 | 60.36 | 29.42 | 10.22 | 50.01 | 1.60 |
| IDT -M | 340.85 | 7.39 | 57.92 | 99.81 | 99.64 | 98.84 | 92.37 | 79.64 | 97.58 | 96.72 | 48.95 | 40.63 | 10.43 | 40.56 | 1.95 |
| IDT -L | 423.60 | 8.60 | 54.28 | 99.80 | 99.32 | 97.01 | 85.53 | 70.18 | 97.39 | 94.83 | 45.15 | 44.10 | 10.75 | 37.41 | 2.28 |
| Roche-S | 258.64 | 11.81 | 60.10 | 99.86 | 99.36 | 98.33 | 93.53 | 83.01 | 96.17 | 95.02 | 51.86 | 18.13 | 30.01 | 35.50 | 1.89 |
| Roche-M | 355.99 | 10.60 | 55.76 | 99.83 | 99.27 | 98.03 | 91.86 | 79.02 | 96.90 | 95.53 | 49.33 | 38.52 | 12.14 | 33.77 | 1.90 |
| Roche-L | 480.35 | 8.75 | 50.31 | 99.80 | 99.11 | 97.37 | 87.91 | 71.05 | 96.79 | 94.84 | 42.28 | 36.04 | 21.68 | 28.94 | 2.02 |
| Agilent-S | 267.89 | 14.89 | 70.25 | 99.79 | 99.33 | 98.36 | 94.21 | 85.84 | 95.23 | 94.22 | 61.57 | 21.20 | 17.23 | 32.85 | 2.04 |
| Agilent-M | 353.80 | 10.88 | 66.15 | 99.75 | 99.33 | 98.49 | 94.48 | 85.58 | 96.27 | 95.40 | 57.04 | 26.17 | 16.79 | 30.44 | 1.96 |
| Agilent-L | 441.38 | 11.48 | 58.31 | 99.74 | 99.16 | 97.73 | 90.82 | 78.15 | 96.13 | 94.62 | 50.92 | 36.96 | 12.13 | 27.17 | 2.06 |
| Twist-S | 209.67 | 5.10 | 61.80 | 99.86 | 99.78 | 99.33 | 95.85 | 89.38 | 95.51 | 95.06 | 52.23 | 33.16 | 14.61 | 45.77 | 1.59 |
| Twist-M | 368.28 | 5.26 | 46.41 | 99.82 | 99.71 | 99.21 | 94.96 | 83.24 | 96.57 | 96.06 | 38.92 | 45.65 | 15.43 | 34.11 | 1.47 |
| Twist-L | 398.16 | 3.60 | 45.17 | 99.82 | 99.69 | 99.05 | 94.00 | 80.94 | 96.56 | 95.90 | 37.24 | 47.04 | 15.72 | 32.63 | 1.51 |

The entire dataset (Supplementary Table 2) was normalized to a 140 theoretical X-fold coverage on the target design and the results were aggregated by the mean value of the three replicates (Table 1). The average insert sizes in the libraries prepared using the short and the medium fragments were 172–268 and 341–368 bp, respectively, whereas the long DNA fragments were often shorter than expected (398–480 bp). We evaluated the frequency of duplicates and the number of sequenced bases near and off the target obtained using different DNA fragment lengths. Short libraries generated the highest frequency of duplicates in three of the four platforms (12–15%), followed by the long libraries in two of the four platforms (9–12%). As expected, the number of sequenced near-target bases increased in all four platforms when the insert size was larger, whereas the off-target rate did not change in two of the four platforms (IDT and Twist) and declined in the other two (Roche and Agilent). The fold enrichment, which is strongly dependent on both the near-target and off-target rates, was lower in all four platforms with larger DNA fragment sizes. The differences in duplicate and off-target rates caused by different DNA fragment lengths resulted in high variability between the theoretical and mapped coverage values for each combination, as anticipated.

### Influence of DNA fragment size and enrichment uniformity on genotypability

To assess the genotypability of the targets using different DNA fragment lengths, we compared base calling at uniform coverage levels for each platform. We therefore analyzed a set of downsampled BAM files with an average deduplicated X-fold coverage of 80 on each target design (Table 2). The enrichment uniformity, evaluated by applying the FOLD 80 penalty value (the fold over-coverage necessary to raise 80% of bases to the mean coverage level in those targets [25]), was influenced by the increase in DNA fragment size. Compared to the short DNA fragments, longer ones increased the uniformity in one platform (Twist) but reduced it in two others (IDT and Roche), suggesting DNA fragment extension had a platform-specific effect. However, the medium and long DNA fragments achieved higher base calling values in all platforms (96.58– 98.23%) compared to the short DNA fragments (95.52–97.58%).

**Table 2.** The 80X mapped dataset. For each platform and DNA fragment length combination, the 80 mapped X-fold coverage is shown for the target design dataset (mean of the three independent experiments). The columns show the average insert size, percentage of reads marked as duplicates, mapped coverage on the target, percentage of the target covered by at least 1, 5, 10, 20 and 30 reads, percentage of callable bases on the target for standard read depth (>3) and read depth >10, percentage of bases on/near/off target, fold enrichment and FOLD 80 base penalty. DNA fragment lengths: S = short, M = medium, L = long.

| ID | Mapped coverage (X) | %1X | %5X | %10X | %20X | %30X | % PASS | % PASS RD>10 | Fold enrichment | FOLD 80 penalty |
| --- | --- | --- | --- | --- | --- | --- | --- | --- | --- | --- |
| IDT-S | 78.04 | 99.83 | 99.77 | 99.67 | 98.72 | 95.26 | 96.81 | 96.71 | 49.99 | 1.60 |
| IDT-M | 80.00 | 99.82 | 99.72 | 99.45 | 97.01 | 90.60 | 97.62 | 97.32 | 40.49 | 1.93 |
| IDT-L | 80.29 | 99.82 | 99.63 | 98.89 | 93.99 | 85.11 | 97.55 | 96.73 | 37.28 | 2.28 |
| Roche-S | 80.30 | 99.88 | 99.55 | 98.96 | 96.71 | 91.89 | 96.29 | 95.61 | 35.46 | 1.86 |
| Roche-M* | 66.83 | 99.85 | 99.40 | 98.54 | 94.63 | 86.06 | 96.99 | 96.01 | 33.74 | 1.91 |
| Roche-L | 79.52 | 99.85 | 99.49 | 98.83 | 95.99 | 89.65 | 97.04 | 96.30 | 28.82 | 2.01 |
| Agilent-S | 77.73 | 99.65 | 99.15 | 98.25 | 94.79 | 88.17 | 97.58 | 96.58 | 32.79 | 2.05 |
| Agilent-M | 80.01 | 99.64 | 99.30 | 98.76 | 96.26 | 90.63 | 98.23 | 97.64 | 30.29 | 1.95 |
| Agilent-L | 76.42 | 99.66 | 99.27 | 98.55 | 94.99 | 87.51 | 98.20 | 97.39 | 26.82 | 2.05 |
| Twist-S | 80.00 | 99.86 | 99.81 | 99.65 | 97.94 | 94.22 | 95.52 | 95.36 | 45.67 | 1.58 |
| Twist-M | 79.96 | 99.84 | 99.78 | 99.70 | 99.08 | 97.11 | 96.58 | 96.51 | 33.52 | 1.42 |
| Twist-L | 80.05 | 99.84 | 99.78 | 99.69 | 98.93 | 96.66 | 96.58 | 96.50 | 32.17 | 1.45 |

\* The sequencing data available for this combination did not reach 80X mapped coverage.

Given the evident influence of DNA fragment extension on both enrichment uniformity and genotypability, we evaluated the single and combined effects of DNA fragment sizes and enrichment uniformity on base calling at different coverage levels by producing downsampled BAM files (with an average deduplicated X-fold coverage of 10–80) on the corresponding target designs. To assess how enrichment uniformity influenced genotypability at different coverage levels, we compared the two platforms with the best (Twist) and worst (Agilent) enrichment uniformity for the medium DNA fragments (Table 3). This revealed that the highest uniformity at 80X coverage corresponded to the best genotypability both at the standard read depth (PASS, 96.58%) and the minimum read depth of 10 (PASS10, 96.51%). The platform with the best uniformity (Twist) achieved PASS saturation at 60X coverage (96.57%), whereas the Agilent platform did not reach saturation and showed a maximum PASS value of 96.31% at 80X coverage. PASS10 values are more relevant in a clinical context, and the difference between platforms was similar (Twist-M = 95.77% at 40X coverage, and Agilent-M = 95.72% at 80X coverage).

**Table 3.** Downsampled mapped coverage for the platforms showing the best and worst FOLD 80 values at fixed DNA fragment lengths. Parameters were calculated on 10–80X downsampled sets, including mapped coverage on the target, percentage of the target covered by at least 1, 5, 10, 20 and 30 reads, percentage of callable bases on the target for standard read depth (>3) and read depth >10, fold enrichment and FOLD 80 base penalty.

| Mapped coverage (X) | %1X | %5X | %10X | %20X | %30X | % PASS | % PASS RD>10 | Fold enrichment | FOLD 80 penalty |
| --- | --- | --- | --- | --- | --- | --- | --- | --- | --- |
| Twist-M |  |  |  |  |  |  |  |  |  |
| 80 | 99.84 | 99.78 | 99.70 | 99.08 | 97.11 | 96.58 | 96.51 | 33.52 | 1.42 |
| 70 | 99.84 | 99.77 | 99.66 | 98.66 | 95.67 | 96.57 | 96.47 | 33.51 | 1.42 |
| 60 | 99.83 | 99.76 | 99.58 | 97.87 | 93.03 | 96.57 | 96.40 | 33.52 | 1.42 |
| 50 | 99.83 | 99.73 | 99.40 | 96.32 | 87.50 | 96.55 | 96.21 | 33.51 | 1.44 |
| 40 | 99.82 | 99.68 | 98.94 | 92.62 | 74.75 | 96.53 | 95.77 | 33.51 | 1.46 |
| 30 | 99.81 | 99.49 | 97.55 | 81.67 | 47.28 | 96.45 | 94.39 | 33.51 | 1.50 |
| 20 | 99.79 | 98.64 | 91.45 | 47.98 | 11.39 | 96.02 | 88.26 | 33.51 | 1.58 |
| 10 | 99.54 | 89.28 | 49.25 | 3.28 | 0.38 | 90.86 | 46.27 | 33.51 | 1.58 |
| Agilent-M |  |  |  |  |  |  |  |  |  |
| 80 | 99.78 | 99.45 | 98.86 | 96.41 | 90.93 | 96.31 | 95.72 | 30.41 | 1.94 |
| 70 | 99.76 | 99.37 | 98.62 | 95.16 | 87.38 | 96.27 | 95.50 | 30.41 | 1.95 |
| 60 | 99.75 | 99.27 | 98.24 | 93.00 | 81.79 | 96.21 | 95.16 | 30.41 | 1.96 |
| 50 | 99.73 | 99.09 | 97.54 | 89.08 | 73.10 | 96.12 | 94.51 | 30.41 | 1.97 |
| 40 | 99.69 | 98.78 | 96.10 | 81.57 | 59.87 | 95.96 | 93.13 | 30.41 | 1.98 |
| 30 | 99.63 | 98.05 | 92.40 | 67.22 | 40.83 | 95.56 | 89.53 | 30.41 | 2.00 |
| 20 | 99.47 | 95.47 | 80.96 | 41.50 | 17.69 | 94.11 | 78.27 | 30.42 | 2.13 |
| 10 | 98.63 | 80.04 | 43.34 | 8.65 | 1.84 | 84.16 | 41.31 | 30.41 | 2.39 |

Next we assessed the influence of the DNA fragment size on genotypability, focusing on the platform showing the most variable enrichment uniformity. IDT showed a sharp decrease in coverage uniformity (a greater increase in the FOLD 80 penalty) when fragment size was longer than recommended by the manufacturer, jumping from 1.6 (IDT-S) to 2.28 (IDT-L) as shown in Table 4. With regard to coverage levels, IDT-L produced a greater number of over-represented regions (higher %30X values) than IDT-S at 10X and 20X mapped coverage, whereas the proportion of the target region covered by 10 or more reads at higher coverage levels (40X to 80X) was much lower for IDT-L, indicating an uneven distribution of longer reads. In terms of genotypability, we observed overall better PASS values with longer DNA fragments even at coverages as low as 40X. However, the beneficial effect of longer reads on genotypability did not overcome the negative effect on enrichment uniformity, given that IDT-L did not achieve higher PASS10 values than IDT-S.

**Table 4.**
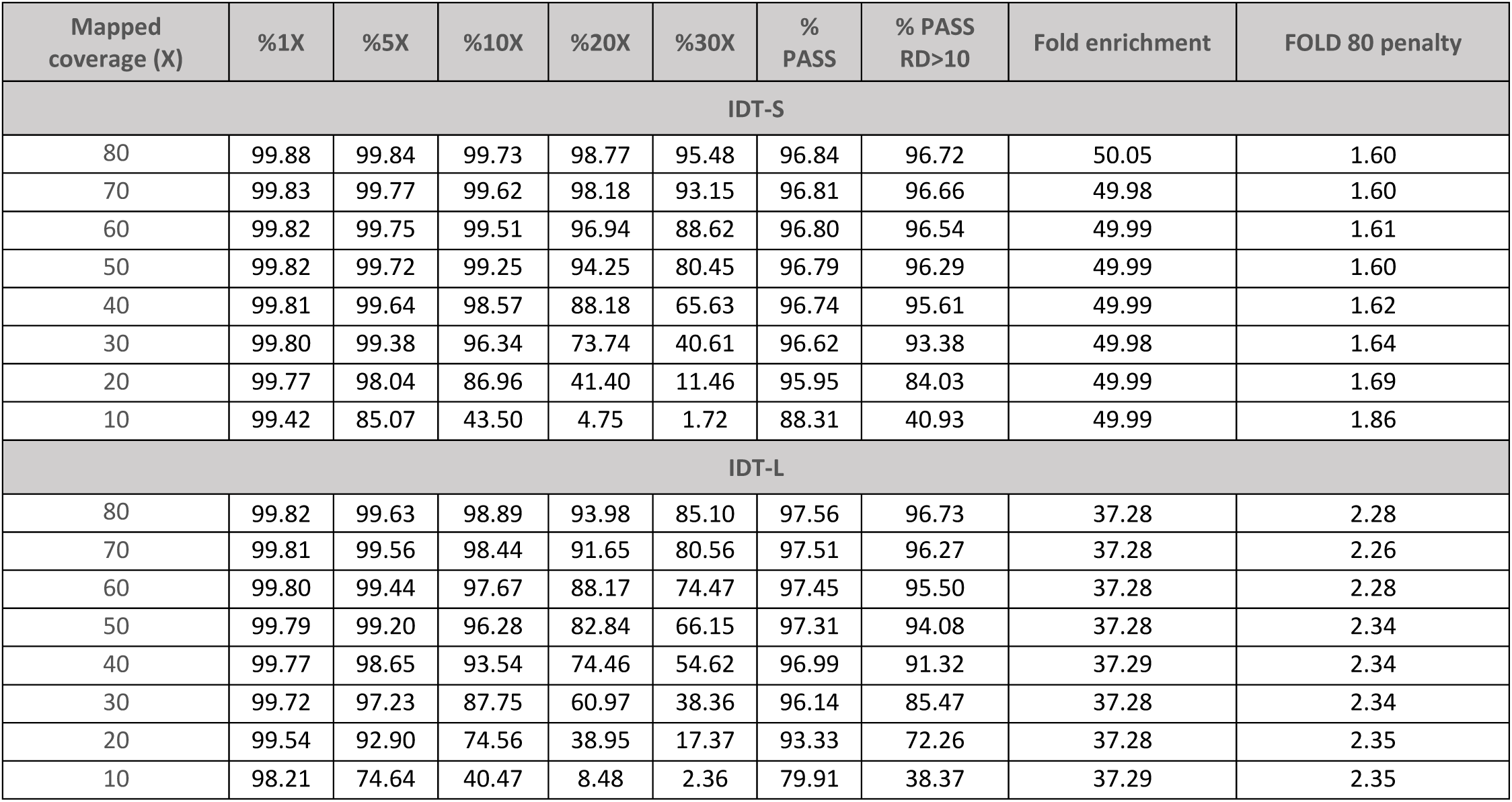
Downsampled mapped coverage for the platform showing the highest variation of FOLD 80 values using different DNA fragment lengths. Parameters were calculated on 10–80X downsampled sets, including mapped coverage on the target, percentage of the target covered by at least 1, 5, 10, 20 and 30 reads, percentage of callable bases on the target for standard read depth (>3) and read depth >10, fold enrichment and FOLD 80 base penalty.

| Mapped coverage (X) | %1X | %5X | %10X | %20X | %30X | % PASS | % PASS RD>10 | Fold enrichment | FOLD 80 penalty |
| --- | --- | --- | --- | --- | --- | --- | --- | --- | --- |
| IDT-S |  |  |  |  |  |  |  |  |  |
| 80 | 99.88 | 99.84 | 99.73 | 98.77 | 95.48 | 96.84 | 96.72 | 50.05 | 1.60 |
| 70 | 99.83 | 99.77 | 99.62 | 98.18 | 93.15 | 96.81 | 96.66 | 49.98 | 1.60 |
| 60 | 99.82 | 99.75 | 99.51 | 96.94 | 88.62 | 96.80 | 96.54 | 49.99 | 1.61 |
| 50 | 99.82 | 99.72 | 99.25 | 94.25 | 80.45 | 96.79 | 96.29 | 49.99 | 1.60 |
| 40 | 99.81 | 99.64 | 98.57 | 88.18 | 65.63 | 96.74 | 95.61 | 49.99 | 1.62 |
| 30 | 99.80 | 99.38 | 96.34 | 73.74 | 40.61 | 96.62 | 93.38 | 49.98 | 1.64 |
| 20 | 99.77 | 98.04 | 86.96 | 41.40 | 11.46 | 95.95 | 84.03 | 49.99 | 1.69 |
| 10 | 99.42 | 85.07 | 43.50 | 4.75 | 1.72 | 88.31 | 40.93 | 49.99 | 1.86 |
| IDT-L |  |  |  |  |  |  |  |  |  |
| 80 | 99.82 | 99.63 | 98.89 | 93.98 | 85.10 | 97.56 | 96.73 | 37.28 | 2.28 |
| 70 | 99.81 | 99.56 | 98.44 | 91.65 | 80.56 | 97.51 | 96.27 | 37.28 | 2.26 |
| 60 | 99.80 | 99.44 | 97.67 | 88.17 | 74.47 | 97.45 | 95.50 | 37.28 | 2.28 |
| 50 | 99.79 | 99.20 | 96.28 | 82.84 | 66.15 | 97.31 | 94.08 | 37.28 | 2.34 |
| 40 | 99.77 | 98.65 | 93.54 | 74.46 | 54.62 | 96.99 | 91.32 | 37.29 | 2.34 |
| 30 | 99.72 | 97.23 | 87.75 | 60.97 | 38.36 | 96.14 | 85.47 | 37.28 | 2.34 |
| 20 | 99.54 | 92.90 | 74.56 | 38.95 | 17.37 | 93.33 | 72.26 | 37.28 | 2.35 |
| 10 | 98.21 | 74.64 | 40.47 | 8.48 | 2.36 | 79.91 | 38.37 | 37.29 | 2.35 |

Finally, we evaluated the combined effect of DNA fragment extension and improved enrichment uniformity on genotypability in the Twist platform, which demonstrated the most beneficial effect of longer DNA fragment sizes on the uniformity of enrichment. The comparison of Twist-S and Twist-M (Table 5) showed that genotypability improved by more than 1% when both the DNA fragment size and the enrichment uniformity increased, and this was the case for both PASS (96.58%) and PASS10 (96.51%) values. DNA fragment extension did not affect PASS saturation, which was reached at 60X for both Twist-M and Twist-S, but resulted in ∼1% more genotyped bases at 80X coverage (96.58% and 95.52%, respectively). The improvement was even higher for the PASS10 values (Twist-M = 96.51%, Twist-S = 95.36%). Therefore, greater enrichment uniformity and medium-length DNA fragments produce a synergistic effect in terms of better genotypability of the target region, especially for clinically-relevant thresholds (PASS10).

**Table 5.** Downsampled mapped coverage for the platform showing the most beneficial effect of longer DNA fragments on FOLD 80 values. Parameters were calculated on 10–80X downsampled sets, including mapped coverage on the target, percentage of the target covered by at least 1, 5, 10, 20 and 30 reads, percentage of callable bases on the target for standard read depth (>3) and read depth >10, fold enrichment and FOLD 80 base penalty.

| Mapped coverage (X) | %1X | %5X | %10X | %20X | %30X | % PASS | % PASS RD>10 | Fold enrichment | FOLD 80 penalty |
| --- | --- | --- | --- | --- | --- | --- | --- | --- | --- |
| Twist-S |  |  |  |  |  |  |  |  |  |
| 80 | 99.86 | 99.81 | 99.65 | 97.94 | 94.22 | 95.52 | 95.36 | 45.67 | 1.58 |
| 70 | 99.86 | 99.79 | 99.52 | 97.02 | 92.09 | 95.51 | 95.25 | 45.68 | 1.58 |
| 60 | 99.86 | 99.77 | 99.28 | 95.56 | 88.66 | 95.50 | 95.01 | 45.67 | 1.60 |
| 50 | 99.85 | 99.71 | 98.80 | 93.13 | 82.65 | 95.48 | 94.53 | 45.67 | 1.60 |
| 40 | 99.84 | 99.56 | 97.74 | 88.39 | 70.86 | 95.41 | 93.50 | 45.67 | 1.62 |
| 30 | 99.83 | 99.09 | 95.29 | 77.32 | 47.59 | 95.18 | 91.09 | 45.67 | 1.66 |
| 20 | 99.78 | 97.36 | 87.52 | 47.91 | 13.60 | 94.22 | 83.48 | 45.67 | 1.71 |
| 10 | 99.30 | 86.10 | 48.97 | 4.46 | 0.59 | 87.35 | 45.76 | 45.67 | 1.88 |
| Twist-M |  |  |  |  |  |  |  |  |  |
| 80 | 99.84 | 99.78 | 99.70 | 99.08 | 97.11 | 96.58 | 96.51 | 33.52 | 1.42 |
| 70 | 99.84 | 99.77 | 99.66 | 98.66 | 95.67 | 96.57 | 96.47 | 33.51 | 1.42 |
| 60 | 99.83 | 99.76 | 99.58 | 97.87 | 93.03 | 96.57 | 96.40 | 33.52 | 1.42 |
| 50 | 99.83 | 99.73 | 99.40 | 96.32 | 87.50 | 96.55 | 96.21 | 33.51 | 1.44 |
| 40 | 99.82 | 99.68 | 98.94 | 92.62 | 74.75 | 96.53 | 95.77 | 33.51 | 1.46 |
| 30 | 99.81 | 99.49 | 97.55 | 81.67 | 47.28 | 96.45 | 94.39 | 33.51 | 1.50 |
| 20 | 99.79 | 98.64 | 91.45 | 47.98 | 11.39 | 96.02 | 88.26 | 33.51 | 1.58 |
| 10 | 99.54 | 89.28 | 49.25 | 3.28 | 0.38 | 90.86 | 46.27 | 33.51 | 1.58 |

### Genotypability of RefSeq and OMIM genes

We also evaluated the genotypability of RefSeq genes using the downsampled BAM files at 80X mapped coverage on the target designs, focusing on the influence of DNA fragment size on the genotypability of each gene. The resulting dataset (Table 6) showed similar trends to those described above. The medium and long DNA fragments achieved higher genotypability (96.54–97.14%) in all platforms compared to the short fragments (95.72–96.28%).

**Table 6.** The 80X mapped dataset (RefSeq genes). For each platform and DNA fragment length combination, the 80 mapped X-fold coverage is shown for the target design dataset (mean of the three independent experiments). The columns show the average insert size, percentage of reads marked as duplicates, mapped coverage on the target, percentage of the target covered by at least 1, 5, 10, 20 and 30 reads, percentage of callable bases on the target for standard read depth (>3) and read depth >10, percentage of bases on/near/off target, fold enrichment and FOLD 80 base penalty. DNA fragment lengths: S = short, M = medium, L = long.

| ID | Mapped coverage (X) | %1X | %5X | %10X | %20X | %30X | % PASS | % PASS RD>10 | Fold enrichment | FOLD 80 penalty |
| --- | --- | --- | --- | --- | --- | --- | --- | --- | --- | --- |
| IDT-S | 78.43 | 99.16 | 99.02 | 98.93 | 98.15 | 94.96 | 95.72 | 95.63 | 50.41 | 1.60 |
| IDT-M | 79.36 | 99.22 | 99.03 | 98.73 | 96.21 | 89.55 | 96.59 | 96.27 | 40.31 | 1.95 |
| IDT-L | 79.61 | 99.26 | 98.96 | 98.17 | 93.07 | 83.84 | 96.54 | 95.68 | 37.10 | 2.31 |
| Roche-S | 82.94 | 99.80 | 99.48 | 98.99 | 97.31 | 93.82 | 96.21 | 95.67 | 36.62 | 1.74 |
| Roche-M* | 69.35 | 99.77 | 99.36 | 98.69 | 95.77 | 89.05 | 96.98 | 96.23 | 35.01 | 1.81 |
| Roche-L | 83.25 | 99.79 | 99.44 | 98.93 | 96.82 | 91.89 | 97.03 | 96.47 | 30.18 | 1.92 |
| Agilent-S | 86.86 | 99.76 | 99.44 | 98.91 | 96.60 | 91.58 | 96.28 | 95.73 | 36.63 | 2.01 |
| Agilent-M | 89.27 | 99.72 | 99.39 | 98.97 | 97.20 | 93.02 | 97.03 | 96.60 | 33.80 | 1.94 |
| Agilent-L | 86.23 | 99.74 | 99.37 | 98.79 | 96.06 | 90.20 | 96.99 | 96.37 | 30.26 | 2.08 |
| Twist-S | 79.50 | 99.81 | 99.75 | 99.60 | 97.90 | 94.18 | 96.18 | 96.04 | 45.39 | 1.58 |
| Twist-M | 79.96 | 99.80 | 99.73 | 99.67 | 99.09 | 97.19 | 97.13 | 97.07 | 33.51 | 1.41 |
| Twist-L | 80.08 | 99.81 | 99.73 | 99.65 | 98.94 | 96.74 | 97.14 | 97.06 | 32.18 | 1.44 |

\* The sequencing data available for this combination did not reach 80X mapped coverage.

We then calculated the number of genes that could reach (i) 100% genotypability and (ii) any increase in genotypability as a consequence of the increase of the DNA fragment size. The medium-size DNA fragments performed best in three of the four platforms, with long libraries performing best in the Roche platform. There was a difference of 1656 genes between the best (Twist-M) and worst (Agilent-S) performing platforms (Table 7). In the first calculation, the platform with the best enrichment uniformity (Twist) reached 100% genotypability for 1107 genes by extending the DNA fragment length (Table 8), including 100 genes that improved by more than 25% in both the short-to-medium and short-to-long fragment extensions (Supplementary Figure 1a). In the second calculation, 1993 genes showed an increase in genotypability (Table 8) due to DNA fragment extension (short-to-medium or short-to-long) including almost 200 genes that improved by more than 30% (Supplementary Figure 1a). Overall, more than 800 RefSeq genes reached 100% genotypability in all four platforms and more than 1800 genes showed some increase in genotypability (Table 8 and Figure 2). Only for a minimal number of the genes which could reach 100% genotypability with the short-size DNA fragments in each platform (99–312), the extension of the DNA fragment length caused a decrease in genotypability (Figure 2).

**Table 7.**
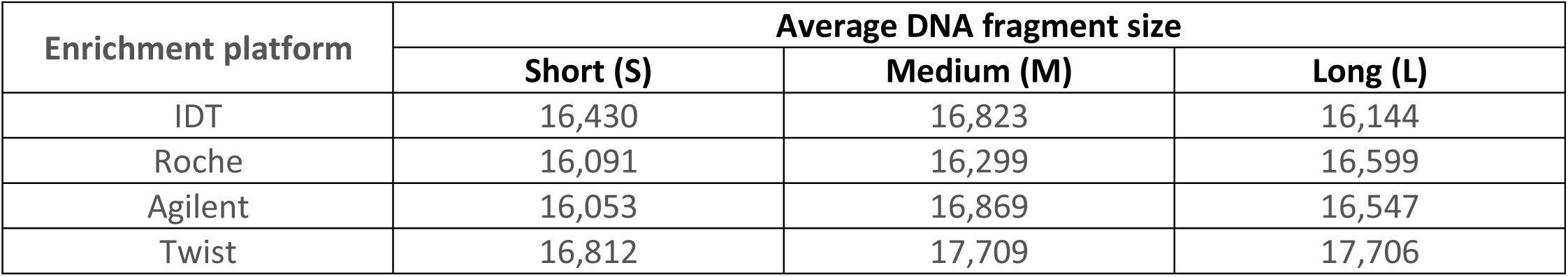
Number of RefSeq genes reaching 100% genotypability. Number of RefSeq genes reaching 100% genotypability at 80X mapped coverage on the target design dataset using different platforms and DNA fragment lengths.

**Table 8.** Number of genes showing increased genotypability. Number of RefSeq and OMIM genes showing increased genotypability following the extension of the DNA fragment size from short to medium, or short to long, at 80X mapped coverage on each target design.

| **Dataset** | **IDT** | **Roche** | **Agilent** | **Twist** |
| --- | --- | --- | --- | --- |
| **RefSeq genes – up to 100% genotypability** | 840 | 1007 | 1330 | 1107 |
| **RefSeq genes – increased genotypability** | 1837 | 2247 | 2429 | 1993 |
| **OMIM genes – up to 100% genotypability** | 156 | 125 | 270 | 232 |
| **OMIM genes – increased genotypability** | 321 | 288 | 459 | 370 |

**Figure 1.**
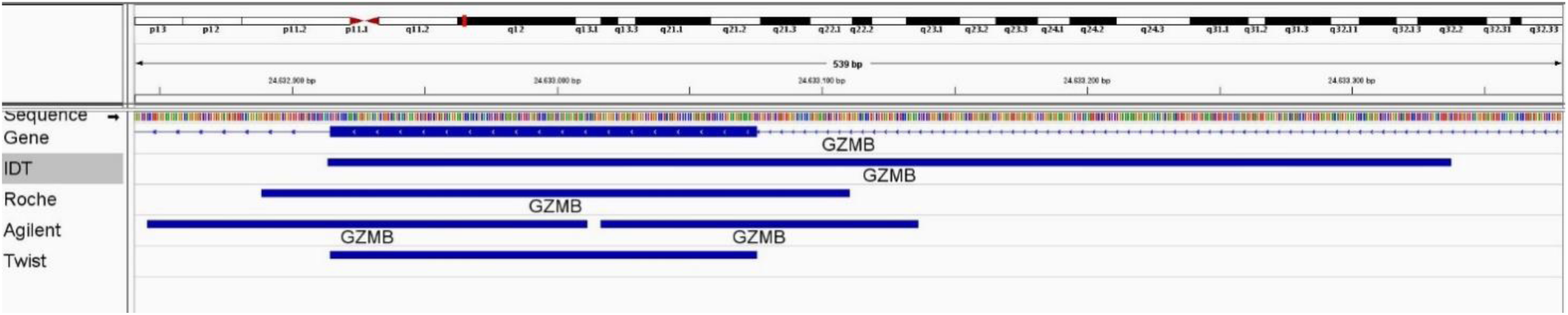
Differences in BED coordinates. Genomic coordinates of exon 2 of the GZMB gene reported in the BED files provided by different enrichment platforms suppliers.

**Figure 2.**
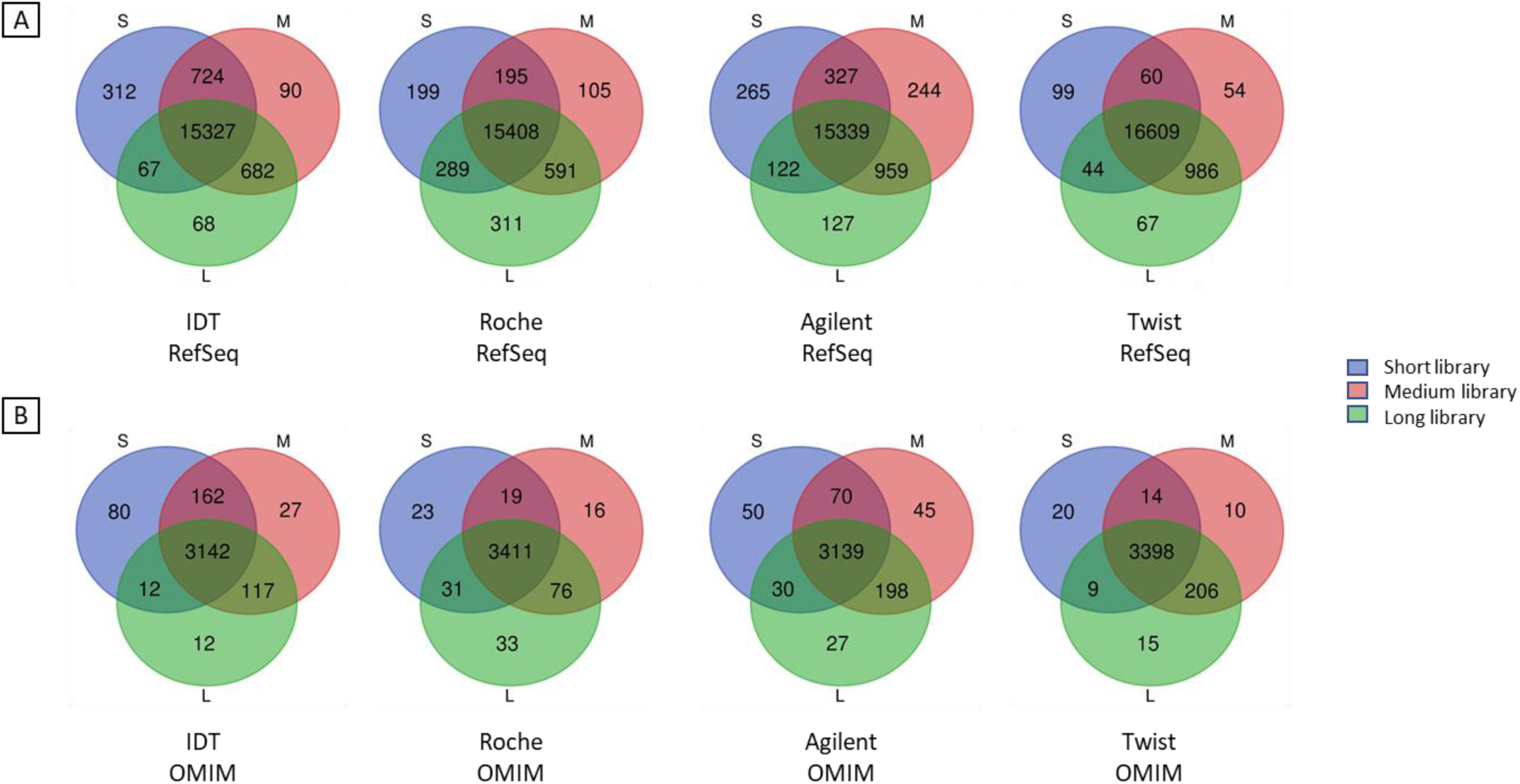
RefSeq/OMIM genes reaching 100% genotypability. Number of RefSeq (A) and OMIM (B) genes reaching 100% genotypability at 80X mapped coverage on each target design using different DNA fragment lengths.

The 3873 OMIM genes associated with a clinical phenotype were analyzed as above to determine the improvement in genotypability achieved with longer DNA fragments (Table 8, Figure 2 and Supplementary Figure 1b). More than 150 OMIM genes reached 100% genotypability in all four platforms, and more than 280 showed some increase in genotypability. The top 20 OMIM genes ranked by improvement in genotypability as a consequence of extended DNA fragment size showed that, at equal coverage levels, the genotypability of the target region increased with DNA fragment length, from 18% to 52% (Table 9). As seen for the RefSeq dataset, a subset of the OMIM genes which could reach 100% genotypability with the short-size DNA fragments (20–80) decreased in genotypability extending the DNA fragment length (Figure 2).

**Table 9.**
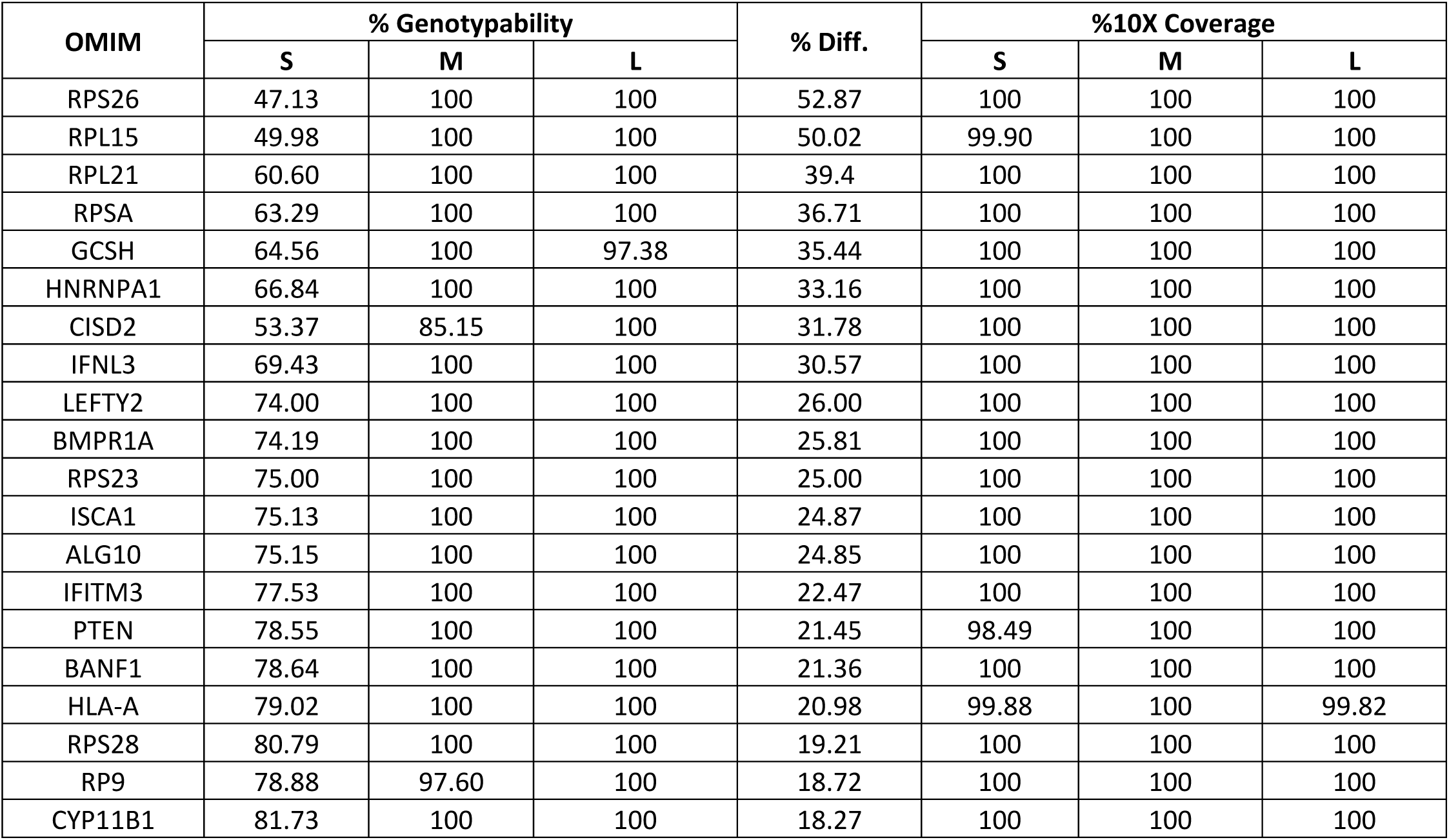
Top 20 OMIM genes showing the best improvement in genotypability. Top 20 OMIM genes showing the best improvement in genotypability following the extension of the DNA fragment length from short to medium and short to long (Twist enrichment platform). The data represent the maximum difference in genotypability at 80X mapped coverage on the Twist design. DNA fragment lengths: S = short, M = medium, L = long.

Finally, for each sample we determined the number of variants present in the Twist target design, the platform showing the most beneficial effect of longer DNA fragments on the uniformity of enrichment. We observed an aggregate mean increase of >1% in both the short-to-medium and short-to-long fragment extensions (Table 10). The same >1% increase with longer DNA fragments was observed for the number of variants identified in the RefSeq and OMIM genes included in the target design. These results reflect the 1% increase in genotypability achieved by increasing the length of the DNA fragments.

**Table 10.** Variants in the Twist target design. Total number of variants identified in the Twist target design, and in the corresponding RefSeq and OMIM genes, for each DNA fragment size (S = short, M = medium, L = long).

| DNA fragment size | #variants in design | #variants in RefSeq genes | #variants in OMIM Genes |
| --- | --- | --- | --- |
| S | 23,140 | 20,291 | 4986 |
| M | 23,461 | 20,529 | 5035 |
| L | 23,521 | 20,599 | 5055 |

## Discussion

Depth of coverage is the parameter used most often to evaluate the performance of WES enrichment technologies, which are applied during the NGS of selected target regions in the genome [7]. However, the genotypability (base calling) of the target provides more comprehensive information, taking into account not only the depth of coverage, but also the quality of the read alignments. We evaluated changes in the genotypability of target regions caused by increasing the DNA fragment size beyond the typical length of the average exon (aimed to reduce the near-target rate) to improve the alignment of reads derived from repetitious genomic regions.

We found that longer DNA inserts increased the mapping quality of reads and thus the mappability of the target region in all four enrichment platforms, suggesting that improvements in base calling can be achieved in these platforms by introducing a measure which is more informative than the standard depth of coverage value. This was particularly evident when evaluating the genotypability of the coding sequence of RefSeq genes at fixed coverage levels (80X mapped). We observed substantial improvements in base calling for many genes, including those of clinical interest in the OMIM dataset. For example, the genotypability of genes *RPL15* and *RPS26* (associated with the bone marrow disorder Diamond-Blackfan anemia) improved to 100%, from 49.98% and 47.13%, respectively. Similarly, the genotypability of *RPSA* (associated with the immunodeficiency disease isolated congenital asplenia) improved from 63.29% to 100%, and that of the tumor suppressor gene *PTEN* improved from 78.55% to 100% (Table 9 and Supplementary Table 3). Longer DNA fragments overcame some of the challenges posed by repetitious genome segments, as recognized by the American College of Medical Genetics and Genomics (ACMG) in their guidelines, which recommend the development of “a strategy for detecting disease-causing variants within regions with known homology” [26]. Expensive long-read sequencing solutions could be adopted for genes that cannot be characterized by short-read sequencing [18] but such methods have yet to be implemented in diagnostic laboratories [27]. Therefore, our new approach offers an alternative solution for the analysis of genes whose read mapping quality is low, although putative pathogenic variants may be present, especially in genes with the highest medical relevance [28].

The number of variants identified among the RefSeq and OMIM genes included in the Twist design showed that it is possible to improve variant calling by increasing the DNA fragment length (Table 10). The 1% increase in genotypability reported above corresponded to an increase of 1% in variant calling, leading to the identification of variants in regions previously considered uncallable because of low mapping quality (Figure 3). The greater number of variants confirms that genotypability is a better parameter for the assessment of WES and that greater mappability corresponds to a higher number of detected variants in repetitious genome regions. Such findings could be clinically significant, especially when analyzing affected patients, and hence they should be included in subsequent variant annotations to prioritize their characterization and assessment.

**Figure 3.**
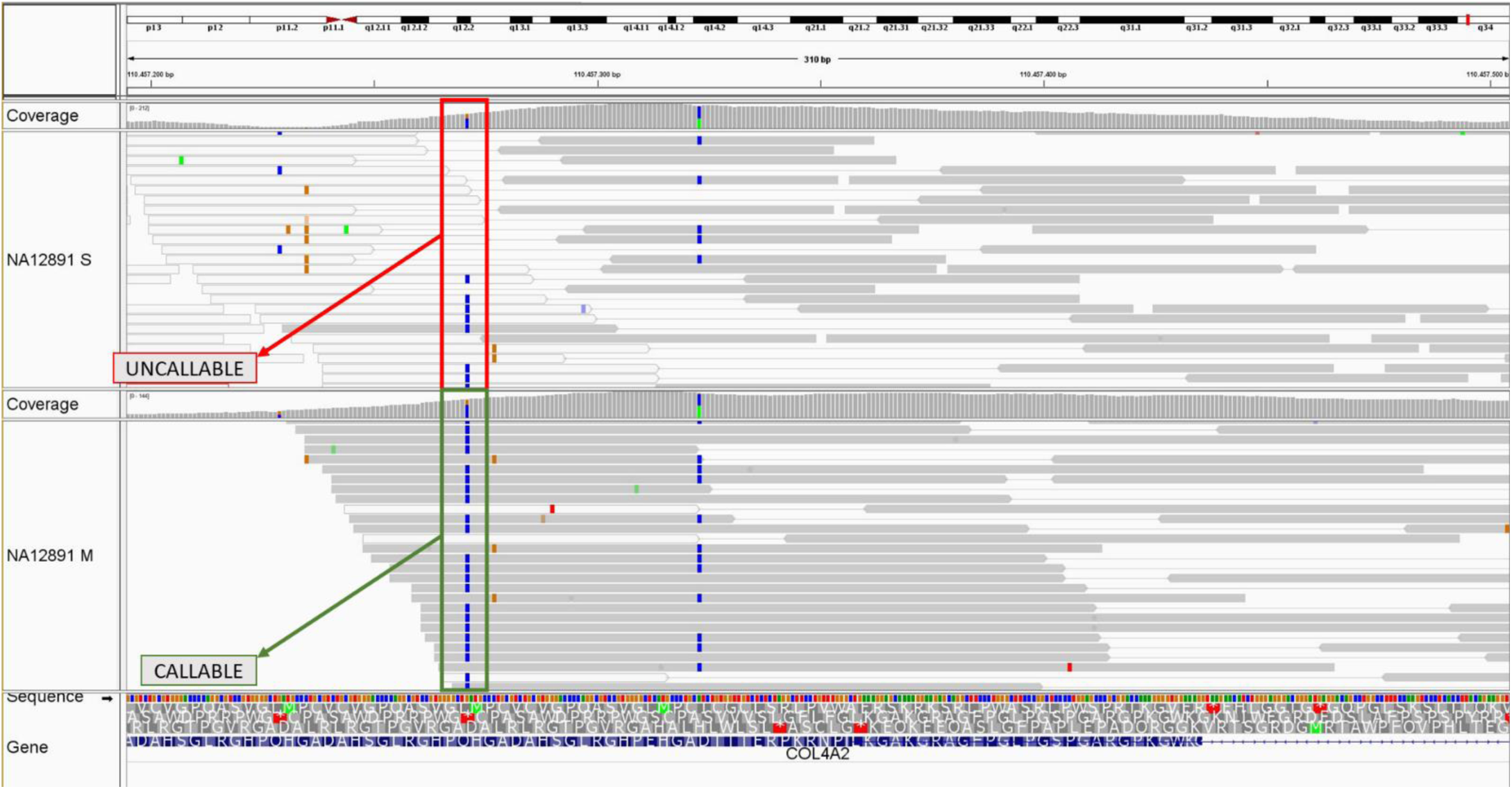
Variant “chr13:110457271” in COL4A2 gene (RefSeq dataset). Variant called in the NA12891 sample using short-size DNA fragment (above) and medium-size DNA fragment (below). The BAM files of the samples are shown on the Twist design at 80X mapped coverage. The colour of the bar indicates the mapping quality of the read: grey = high quality mapping; white = low quality mapping.

Longer DNA inserts do not always improve the uniformity of coverage in the target region, as previously reported [6], and this is another important parameter for the evaluation of enrichment efficiency during targeted NGS. Indeed, the four platforms responded differently in terms of enrichment uniformity: whereas the Roche and Agilent platforms were largely insensitive to the extension of DNA fragment length, the IDT platform showed a dramatic increase in the FOLD 80 penalty (corresponding to low enrichment uniformity), indicating that it has already been optimized for very short fragments. Interestingly, the opposite trend was apparent in the Twist platform, indicating that the already highly uniform enrichment can be improved even further. It would be interesting to determine whether this reflects the internal calibration procedure of the Twist platform or a favorable effect of the double-stranded capture probes. Generally, with high FOLD 80 penalty scores, the genotypability of the target region was directly related to the mapped coverage (higher coverage = higher genotypability), whereas higher enrichment uniformity resulted in the genotypability reaching a plateau at ∼60X coverage, making deeper coverage unnecessary. The advantages of higher enrichment uniformity include the reduction of WES costs and the need for less starting material, given that reducing the depth of sequencing also reduces the number of duplicates. Moreover, increasing the DNA fragment length also helps to overcome the problem of duplicates because for short DNA inserts the use of 2 × 75 bp reads requires the sequencing of twice as many fragments from the same amount of DNA compared to 2 × 150 bp reads. In contrast, the fold enrichment value (another important measure of enrichment efficiency based on the on-target and off-target rates) was often misleading for two reasons. First, fold enrichment depends on the definition of the target region, which in some cases is delineated by the exon boundaries but in others corresponds to a much broader area (Figure 1). Second, fold enrichment does not provide comprehensive information about the real efficiency of the enrichment platforms, given that lower values did not correlate with a reduction in genotypability (Table 2 and Table 6).

Taken together, our data show that WES performance should be based on the genotypability of the target region, which strictly depends on a combination of the DNA fragment size and the uniformity of the enrichment platform. This parameter will help clinicians to select the optimal combination of DNA insert length and enrichment platform during the design of the target region, allowing the correct interpretation of truly positive, but especially truly negative, findings. Our new approach will help to overcome current challenges caused by the presence of repetitious regions in the human genome.

## Acknowledgements

We gratefully acknowledge the Centro Piattaforme Tecnologiche (CPT) for granting access to the genomic facility of University of Verona, and MSc student Andrei Florea for contributing to the bioinformatics pipeline implemented for sequence data analysis.

