## Supplementary Materials for "Enhanced targeted resequencing by optimizing the combination of enrichment technology and DNA fragment length"

*Supplementary Table 1. Library preparation specifications for each enrichment platform.*

| ENRICHMENT PLATFORM | INPUT (ng) | INSERT SIZE (bp) | DNA shearing | | AMPure size selection | PCR cycles |
|---|---|---|---|---|---|---|
| | | | COVARIS treatment time (sec) | Enzymatic digestion time (min) | | |
| IDT | 100 | 150 | 375 | | | 10 |
| | | 350 | 110 | | 0.75X | 10 |
| | | 500 | 85 | | 0.60X | 12 |
| ROCHE | 500 | 200 | 235 | | | 5 |
| | | 350 | 110 | | 0.75X | 5 |
| | | 500 | 85 | | 0.60X | 7 |
| AGILENT | 1500 | 200 | 235 | | | 6 |
| | | 350 | 110 | | 0.75X | 6 |
| | | 500 | 85 | | 0.60X | 8 |
| TWIST | 50 | 200 | | 22 | | 10 |
| | | 350 | 110 | | 0.7X | 10 |
| | | 500 | 85 | | 0.65X | 10 |

*Supplementary Table 1. **Library preparation specifications for each enrichment platform**.*

*Supplementary Table 2. WES initial dataset.*

| ID | Sequenced fragments | GC% | Design length | Theoretical coverage (X) | Mapped deduplicated reads | Average insert size | % Duplicates | Mapped coverage (X) |
|---|---|---|---|---|---|---|---|---|
| NA12891_IDT-S | 41,735,851 | 52 | 38,871,205 | 161.05 | 34,798,458 | 170.56 | 14.76 | 78.02 |
| NA12891_IDT-M | 31,721,147 | 51 | 38,871,205 | 244.82 | 27,295,886 | 338.36 | 12.57 | 95.92 |
| NA12891_IDT-L | 35,056,454 | 52 | 38,871,205 | 270.56 | 29,048,970 | 419.16 | 15.62 | 97.28 |
| NA12891_Roche-S | 69,139,041 | 48 | 47,007,710 | 220.62 | 53,971,159 | 250.20 | 19.09 | 92.04 |
| NA12891_Roche-M | 30,139,859 | 48 | 47,007,710 | 192.35 | 25,187,948 | 352.58 | 14.76 | 73.15 |
| NA12891_Roche-L | 37,597,648 | 48 | 47,007,710 | 239.95 | 31,408,182 | 475.51 | 14.05 | 81.22 |
| NA12891_Agilent-S | 69,997,150 | 51 | 60,448,148 | 173.70 | 56,394,632 | 267.56 | 16.11 | 85.70 |
| NA12891_Agilent-M | 43,377,376 | 50 | 60,448,148 | 215.28 | 36,041,040 | 350.30 | 15.74 | 96.23 |
| NA12891_Agilent-L | 62,476,876 | 49 | 60,448,148 | 310.07 | 50,712,887 | 438.63 | 15.70 | 122.77 |
| NA12891_Twist-S | 62,145,209 | 52 | 36,715,240 | 253.89 | 53,990,842 | 211.43 | 9.24 | 106.91 |
| NA12891_Twist-M | 45,101,242 | 49 | 36,715,240 | 368.52 | 37,418,907 | 390.49 | 14.57 | 106.47 |
| NA12891_Twist-L | 56,814,853 | 49 | 36,715,240 | 464.23 | 48,349,215 | 389.09 | 12.24 | 136.39 |
| NA12892_IDT-S | 39,279,677 | 52 | 38,871,205 | 151.58 | 33,418,710 | 174.23 | 12.97 | 74.92 |
| NA12892_IDT-M | 29,737,235 | 51 | 38,871,205 | 229.51 | 25,829,978 | 340.75 | 11.40 | 90.66 |
| NA12892_IDT-L | 31,442,649 | 52 | 38,871,205 | 242.67 | 26,557,527 | 422.28 | 13.69 | 88.66 |
| NA12892_Roche-S | 63,475,661 | 48 | 47,007,710 | 202.55 | 51,637,279 | 263.49 | 15.33 | 86.49 |
| NA12892_Roche-M | 25,113,466 | 48 | 47,007,710 | 160.27 | 21,737,831 | 353.95 | 11.70 | 63.42 |
| NA12892_Roche-L | 35,647,295 | 48 | 47,007,710 | 227.50 | 29,755,564 | 483.46 | 13.90 | 76.69 |
| NA12892_Agilent-S | 63,367,878 | 50 | 60,448,148 | 157.25 | 50,458,064 | 270.99 | 16.64 | 76.08 |
| NA12892_Agilent-M | 38,379,719 | 50 | 60,448,148 | 190.48 | 32,736,504 | 357.00 | 13.32 | 87.30 |
| NA12892_Agilent-L | 59,863,365 | 49 | 60,448,148 | 297.10 | 49,779,709 | 446.38 | 13.80 | 117.87 |
| NA12892_Twist-S | 58,556,655 | 52 | 36,715,240 | 239.23 | 51,774,402 | 207.55 | 7.76 | 102.74 |
| NA12892_Twist-M | 53,142,446 | 49 | 36,715,240 | 434.23 | 44,835,844 | 360.36 | 13.57 | 131.09 |
| NA12892_Twist-L | 53,633,893 | 49 | 36,715,240 | 438.24 | 46,784,910 | 411.80 | 9.39 | 130.16 |
| VR00_IDT-S | 43,858,514 | 52 | 38,871,205 | 169.25 | 36,430,811 | 170.05 | 14.88 | 81.28 |
| VR00_IDT-M | 30,647,544 | 51 | 38,871,205 | 236.53 | 26,574,858 | 343.45 | 11.82 | 92.66 |
| VR00_IDT-L | 28,719,060 | 51 | 38,871,205 | 221.65 | 24,489,849 | 429.35 | 13.02 | 80.87 |
| VR00_Roche-S | 69,341,641 | 47 | 47,007,710 | 221.27 | 55,349,772 | 262.24 | 16.77 | 80.89 |
| VR00_Roche-M | 25,769,903 | 48 | 47,007,710 | 164.46 | 22,210,538 | 361.45 | 11.86 | 64.00 |
| VR00_Roche-L | 37,393,464 | 47 | 47,007,710 | 238.64 | 31,375,345 | 482.07 | 13.60 | 80.81 |
| VR00_Agilent-S | 63,423,698 | 51 | 60,448,148 | 157.38 | 51,058,660 | 265.11 | 17.83 | 77.59 |
| VR00_Agilent-M | 37,984,977 | 50 | 60,448,148 | 188.52 | 31,756,515 | 354.10 | 15.15 | 84.83 |
| VR00_Agilent-L | 34,450,041 | 49 | 60,448,148 | 170.97 | 28,964,914 | 439.13 | 13.13 | 69.93 |
| VR00_Twist-S | 58,092,508 | 52 | 36,715,240 | 237.34 | 51,158,376 | 209.79 | 8.01 | 101.11 |
| VR00_Twist-M | 49,286,947 | 49 | 36,715,240 | 402.72 | 42,067,556 | 360.41 | 12.36 | 121.92 |
| VR00_Twist-L | 53,025,771 | 48 | 36,715,240 | 433.27 | 46,216,132 | 400.91 | 9.92 | 128.25 |

*Supplementary Table 2. **WES initial dataset**.*
*For each replicate, platform and DNA fragment length combination, the number of sequenced fragments, percentage GC content, theoretical coverage, number of mapped reads without duplicates, average insert size, percentage of reads marked as duplicates and mapped coverage on the target are shown. DNA fragment lengths: S = short, M = medium, L = long.*

| Platform | Short (% PASS) | Medium (% PASS) | Long (% PASS) | % Diff. |
|---|---|---|---|---|
| **RPL15** | | | | |
| IDT | 34.49 | 88.74 | 88.74 | 54.25 |
| Roche | 84.90 | 100.00 | 100.00 | 15.10 |
| Agilent | 83.50 | 100.00 | 100.00 | 16.50 |
| Twist | 49.98 | 100.00 | 100.00 | 50.02 |
| **RPS26** | | | | |
| IDT | 69.35 | 100.00 | 100.00 | 30.65 |
| Roche | 92.05 | 100.00 | 100.00 | 7.95 |
| Agilent | 93.39 | 100.00 | 100.00 | 6.61 |
| Twist | 47.13 | 100.00 | 100.00 | 52.87 |
| **RPSA** | | | | |
| IDT | 69.45 | 100.00 | 100.00 | 30.55 |
| Roche | 96.25 | 100.00 | 100.00 | 3.75 |
| Agilent | 95.46 | 100.00 | 100.00 | 4.54 |
| Twist | 63.29 | 100.00 | 100.00 | 36.71 |
| **PTEN** | | | | |
| IDT | 77.58 | 100.00 | 100.00 | 22.42 |
| Roche | 84.63 | 100.00 | 100.00 | 15.37 |
| Agilent | 90.89 | 100.00 | 100.00 | 9.11 |
| Twist | 78.55 | 100.00 | 100.00 | 21.45 |

*Supplementary Table 3.* ***Genotypability of RPL15, RPS26, RPSA and PTEN genes***

*Supplementary Figure 1a. Increase in the genotypability of RefSeq genes for Twist using different DNA fragment lengths.*

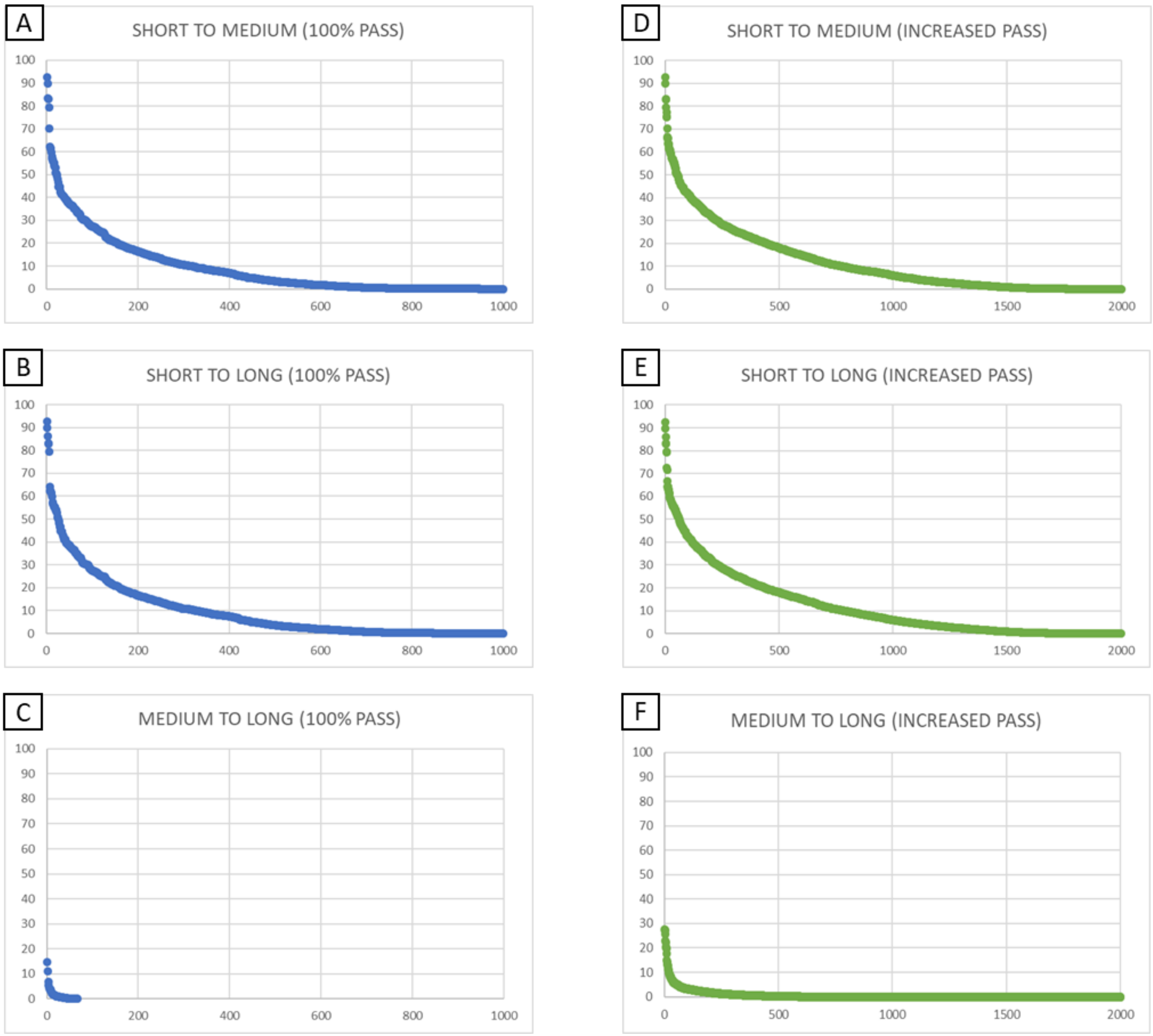


*Supplementary Figure 1a.* ***Increase in the genotypability of RefSeq genes for Twist using different DNA fragment lengths.***

*The percentage increase in genotypability of RefSeq genes at 80X mapped coverage is shown on the Twist design, considering the DNA fragment extension from short to medium (A-D), from short to long (B-E) and from medium to long (C-F) for the genes that reached 100% genotypability (A-B-C) and those showing any increase in genotypability due to an extension of the DNA fragment size (D-E-F).*

*Supplementary Figure 1b. Increase in the genotypability of OMIM genes for Twist using different DNA fragment lengths.*

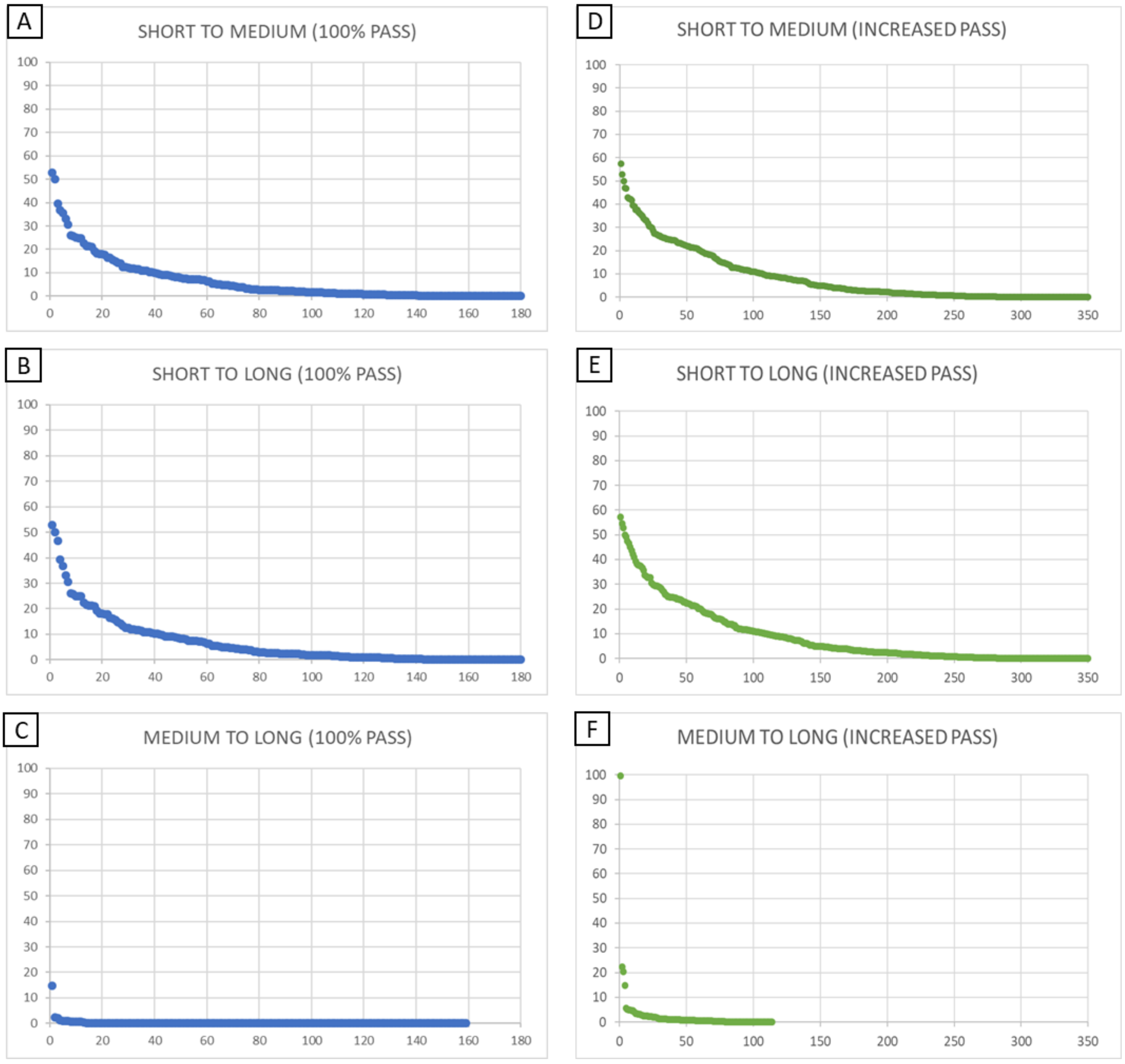


*Supplementary Figure 1b.* ***Increase in the genotypability of OMIM genes for Twist using different DNA fragment lengths***

*The percentage increase in genotypability of OMIM genes at 80X mapped coverage is shown on the Twist design, considering the DNA fragment extension from short to medium (A-D), from short to long (B-E) and from medium to long (C-F) for the genes that reached 100% genotypability (A-B-C) and those showing any increase in genotypability due to an extension of the DNA fragment size (D-E-F).*
